# 6-gingerol interferes with amyloid-beta (Aβ) peptide aggregation

**DOI:** 10.1101/2021.01.03.425159

**Authors:** Elina Berntsson, Suman Paul, Sabrina B. Sholts, Jüri Jarvet, Andreas Barth, Astrid Gräslund, Sebastian K. T. S. Wärmländer

## Abstract

Alzheimer’s disease (AD) is the most prevalent age-related cause of dementia. AD affects millions of people worldwide, and to date there is no cure. The pathological hallmark of AD brains is deposition of amyloid plaques, which mainly consist of amyloid-β (Aβ) peptides, commonly 40 or 42 residues long, that have aggregated into amyloid fibrils. Intermediate aggregates in the form of soluble Aβ oligomers appear to be highly neurotoxic. Cell and animal studies have previously demonstrated positive effects of the molecule 6-gingerol on AD pathology. Gingerols are the main active constituents of the ginger root, which in many cultures is a traditional nutritional supplement for memory enhancement. Here, we use biophysical experiments to characterize *in vitro* interactions between 6-gingerol and Aβ_40_ peptides. Our experiments with atomic force microscopy imaging, and nuclear magnetic resonance and Thioflavin-T fluorescence spectroscopy, show that the hydrophobic 6-gingerol molecule interferes with formation of Aβ_40_ aggregates, but does not interact with Aβ_40_ monomers. Thus, together with its favourable toxicity profile, 6-gingerol appears to display many of the desired properties of an anti-AD compound.

## INTRODUCTION

Alzheimer’s disease (AD) is a progressive and currently incurable neurodegenerative disorder, and the leading cause of age-related dementia worldwide (Frozza *et al.*, 2018; Querfurth and LaFerla, 2010). Although AD brains typically display signs of neuroinflammation and oxidative stress (Agostinho *et al.*, 2010; Regen *et al.*, 2017; Wang *et al.*, 2014b), the main characteristic lesions in AD brains are extracellular amyloid plaques (Querfurth and LaFerla, 2010; Selkoe and Hardy, 2016), which mainly consist of insoluble fibrillar aggregates of amyloid-β (Aβ) peptides (Querfurth and LaFerla, 2010).

The Aβ peptides comprise 37-43 residues and are intrinsically disordered in aqueous solution. They have limited solubility in water due to the hydrophobicity of the central and C-terminal segments, which may fold into a hairpin conformation upon aggregation (Abelein *et al.*, 2014; Baronio *et al.*, 2019). The charged N-terminal segment of Aβ peptides is hydrophilic and interacts readily with cationic molecules and metal ions (Luo *et al.*, 2014a; Owen *et al.*, 2019; Wärmländer *et al.*, 2013).

The Aβ fibrils and plaques that characterize AD neuropathology are the end-products of Aβ aggregation processes (Owen *et al.*, 2019; Selkoe and Hardy, 2016) that involve extra- and/or intracellular formation of intermediate, soluble, and likely neurotoxic Aβ oligomers (Luo *et al.*, 2014b; Sengupta *et al.*, 2016) which may transfer from neuron to neuron via e.g. exosomes (Sardar Sinha *et al.*, 2018). Oligomers of Aβ42 appear to be the most cell-toxic species (Sengupta *et al.*, 2016). The formation of Aβ oligomers is influenced by interactions with various entities such as cellular membranes, small molecules, other proteins, and metal ions (Luo *et al.*, 2016a, b; Owen *et al.*, 2019; Wärmländer *et al.*, 2019; Österlund *et al.*, 2018a). Significant effort has been put into finding suitable molecules - i.e., drug candidates - that may modulate the Aβ aggregation processes (Leshem *et al.*, 2019; Luo *et al.*, 2013; Richman *et al.*, 2013), but so far no drug has been approved (Frozza *et al.*, 2018).

Some investigations of potential anti-AD substances have focused on natural plant compounds, such as gingerols, which are phenolic phytochemical compounds present in the subterranean stem, or rhizome, of angiosperms of the ginger (Zingiberaceae) family (Wang *et al.*, 2014a). Consumed worldwide as a spice and herbal medicine, the rhizome of ginger *(Zingiber officinale)* has demonstrated anti-inflammatory, antioxidant, antiemetic, analgesic, and antimicrobial effects (Sharifi-Rad *et al.*, 2017). Ginger is a common ingredient in traditional healthy diets in many cultures (Iranshahy and Javadi, 2019; Khodaie and Sadeghpoor, 2015). According to Arabian folk wisdom, ginger improves memory and enhances cognition (Saenghong *et al.*, 2012). Gingerols are generally considered to be safe for humans (Kaul and Joshi, 2001; Wang *et al.*, 2014a). Yet, they are cytotoxic towards blood cancer and lung cancer cells (de Lima *et al.*, 2018; Semwal *et al.*, 2015), and *in vitro* studies have demonstrated positive effects also on bowel (Jeong *et al.*, 2009), breast (Lee *et al.*, 2008), ovary (Rhode *et al.*, 2007), and pancreas cancer (Park *et al.*, 2006).

The major pharmacologically-active variant is 6-gingerol, which has been associated with the prevention and treatment of neurodegenerative diseases such as AD (Choi *et al.*, 2018; Jeong *et al.*, 2013; Mohd Sahardi and Makpol, 2019; Wang *et al.*, 2014a). Its chemical structure is shown in Fig. 1. The anti-oxidant and anti-inflammatory properties of 6-gingerol are potentially useful against AD (Mohd Sahardi and Makpol, 2019), which may explain why 6-gingerol has been reported to reduce markers for neuroinflammation and oxidative stress, as well as decrease Aβ levels, in mice and cell AD models (Halawany *et al.*, 2017; Zeng *et al.*, 2015). Little is however known about the molecular mechanisms by which 6-gingerol exerts its positive effects on the AD pathology models. For example, interactions between gingerols and Aβ peptides have not been studied at the molecular level.

**Figure 1.**
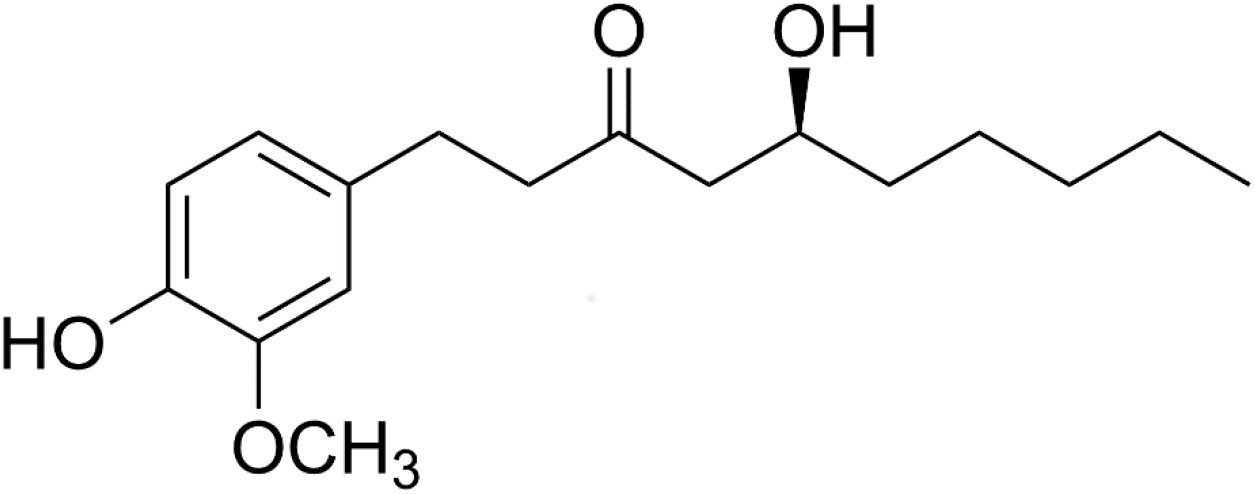
Chemical structure for the hydrophobic plant metabolite 6-gingerol. MW = 294.4 g/mol.

Here, we use biophysical techniques – liquid-phase fluorescence and nuclear magnetic resonance (NMR) spectroscopy together with solid-state atomic force microscopy (AFM) - to investigate possible *in vitro* interactions between 6-gingerol and Aβ_40_ peptides, and how such interactions may affect the Aβ_40_ aggregation and amyloid formation processes.

## MATERIALS AND METHODS

### Reagents and sample preparation

6-gingerol was purchased as a powder from Sigma-Aldrich Inc. (USA), and dissolved in DMSO (dimethyl sulfoxide).

Recombinant unlabeled or uniformly ^15^N-labeled Aβ_40_ peptides, with the primary sequence

DAEFR_5_HDSGY_10_EVHHQ_15_KLVFF_20_AEDVG_25_SNKGA_30_IIGLM_35_VGGVV_40_, were purchased lyophilized from AlexoTech AB (Umeå, Sweden). The peptides were stored at −80 °C until used. The peptide concentration was determined by weight, and the peptide samples were dissolved to monomeric form immediately before each measurement. In brief, the peptides were dissolved in 10 mM sodium hydroxide, pH 12, at a 1 mg/ml concentration and sonicated in an ice-bath for at least three minutes to avoid having pre-formed aggregates in the peptide solutions. The peptide solution was then further diluted in 20 mM buffer of either sodium phosphate or MES (2-[N-morpholino]ethanesulfonic acid) at pH 7.35. All sample preparation steps were performed on ice.

### ThT fluorescence monitoring Aβ aggregation kinetics

To monitor the effect of 6-gingerol on Aβ_40_ aggregation kinetics, 15 μM monomeric Aβ_40_ peptides were incubated in 20 mM MES buffer pH 7.35 in the presence of five different concentrations of 6-gingerol (15, 75, 150, 300, and 1500 μM) together with DMSO (0.1%, 0.6%, 1%, 2% and 10%; vol/vol). Additionally, a control sample without 6-gingerol but containing 2% DMSO was prepared. All samples contained 50 μM Thioflavin T (ThT), which is a benzothiazole dye that displays increased fluorescence intensity when bound to amyloid aggregates (Gade Malmos *et al.*, 2017). The ThT dye was excited at 440 nm, and the fluorescence emission at 480 nm was measured every five minutes in a 96-well plate in a FLUOstar Omega microplate reader (BMG LABTECH, Germany). The sample volume in each well was 35 μl, four replicates per condition were measured, the temperature was +37 °C, and each five-minute cycle involved 140 seconds of shaking at 200 rpm. The assay was repeated three times.

Even though the ThT fluorescence signal reached its maximum value after about seven hours, the incubation in the microplate reader continued for 72 hours to allow the samples to aggregate into mature fibrils that could be observed with AFM imaging (below).

To derive parameters for the aggregation kinetics, the ThT fluorescence curves were fitted to the sigmoidal equation 1:

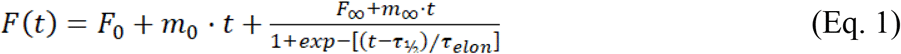

where F_0_ and F_∞_ are the intercepts of the initial and final fluorescence intensity baselines, m_0_ and m_∞_ are the slopes of the initial and final baselines, τ_½_ is the time needed to reach halfway through the elongation phase (i.e., aggregation half-time), and τ_elon_ is the elongation time constant (Gade Malmos *et al.*, 2017). The apparent maximum rate constant for fibrillar growth, r_max_, is defined as 1/τelon.

### Atomic force microscopy (AFM) imaging of Aβ fibrils

Samples for AFM imaging were taken from the samples used in the ThT fluorescence measurements, after 72 h of incubation. AFM images were recorded for the two control samples of 15 μM Aβ_40_ in MES buffer, with and without 2% added DMSO, and for the three samples of 15 μM Aβ_40_ together with 15 μM, 75 μM, and 300 μM of 6-gingerol. Droplets of 1 μl incubated sample were placed on fresh silicon wafers (Siegert Wafer GmbH, Germany) and allowed to sit for 2 minutes. Next, 10 μl Milli-Q water was added to the droplets, and all excess fluid was removed immediately with a lint-free wipe. The wafers were left to dry in a covered container to protect from dust, and AFM images were recorded on the same day. A neaSNOM scattering-type near-field optical instrument (Neaspec GmbH, Germany) was used to collect the AFM images under tapping mode (Ω: 280 kHz, tapping amplitude 50-55 nm) using Pt/Ir-coated monolithic ARROW-NCPt Si tip (NanoAndMore GmbH, Germany) with tip radius <10 nm. Images were acquired on 2.5 x 2.5 μm scan-areas (200 x 200-pixel size) under optimal scan-speed (i.e., 2.5 ms/pixel), and both topographic and mechanical phase images were recorded. Images were minimally processed using the Gwyddion software where a basic plane levelling was performed (Nečas and Klapetek, 2012).

### Nuclear magnetic resonance (NMR) spectroscopy

An Avance 700 MHz NMR spectrometer (Bruker Inc., USA) equipped with a cryogenic probe was used to record 2D ^1^H-^15^N-HSQC spectra at +20 °C of 92.4 μM monomeric ^15^N-labeled Aβ_40_ peptides (500 μl), either in only 20 mM sodium phosphate buffer at pH 7.35 (90/10 H2O/D2O), or in phosphate buffer together with 50 mM SDS (sodium dodecyl sulphate) detergent. As the critical micelle concentration (CMC) for SDS is around 8 mM (Österlund *et al.*, 2018b), most of the SDS was present as micelles. Both samples were titrated, first with additions of pure DMSO, and then by 6-gingerol dissolved in DMSO. The NMR data was processed with the Topspin version 3.6.2 software, and the Aβ_40_ HSQC crosspeak assignment in buffer (Danielsson *et al.*, 2006) and in SDS micelles (Jarvet *et al.*, 2007) is known from previous work.

## RESULTS

### ThT fluorescence kinetics

Fig. 2 shows ThT fluorescence intensity curves for 15 μM Aβ_40_ peptides, incubated in the presence of varying concentrations of 6-gingerol and DMSO. These curves reflect the formation of amyloid aggregates, and they all display a generally sigmoidal shape. Fitting Eq. 1 to the curves produces the kinetic parameters τ_½_, rmax, and τ_lag_ (Table 1). Addition of DMSO alone, which was used to dissolve the 6-gingerol, has minor effects on the aggregation kinetics, i.e. by slightly increasing the lag time from 0.94 to 0.98 hrs and decreasing the aggregation half time from 2.2 to 1.9 hrs (Fig. 2, Table 1). With 6-gingerol, some additions produce aggregation kinetics that differ from the control samples. For example, addition of 75 μM 6-gingerol appears to slow down the aggregation (τ_lag_ = 1.3 h; τ_½_ = 3.3 h), while addition of 150 μM 6-gingerol appears to speed up the aggregation (τ_lag_ = 0.5 h; τ_½_ = 1.7 h). There is however variation in these measurements, and there is no overall trend of faster or slower kinetics for the series of 6-gingerol additions. Thus, these data indicate that 6-gingerol has no systematic effect on Aβ_40_ aggregation or amyloid formation.

**Figure 2.**
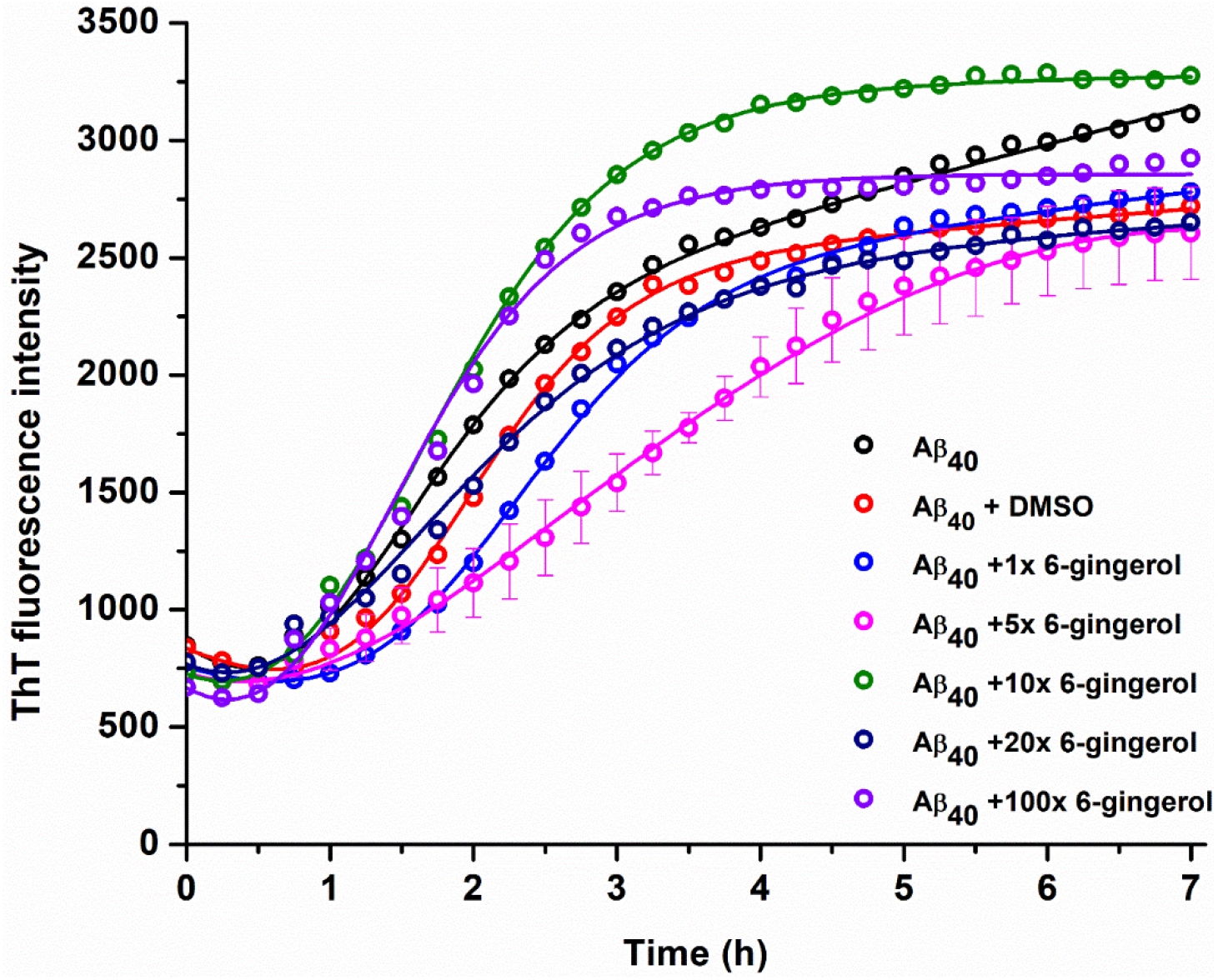
ThT fluorescence curves showing the aggregation kinetics of 15 μM Aβ_40_ in 20 mM MES buffer, pH 7.35, at 37 °C. Black: buffer only; Red: 2% DMSO; Blue: 15 μM 6-gingerol; Pink: 75 μM 6-gingerol; Green: 150 μM 6-gingerol; Dark blue: 300 μM 6-gingerol; and Purple: 1500 μM 6-gingerol. Average curves from four replicates are shown.

**Table 1.**
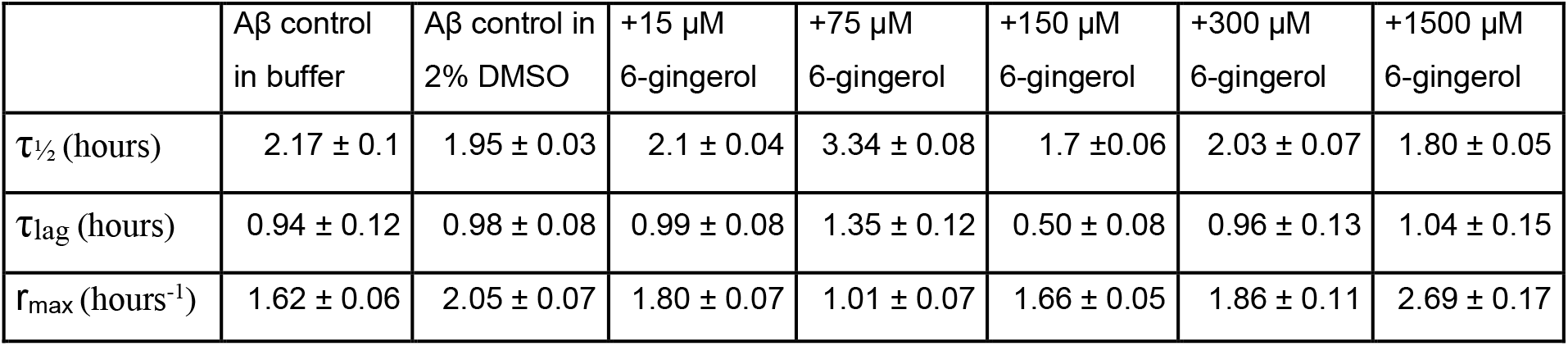
Kinetic parameters (τ_½_, τ_lag_, and Γ_max_) for fibril formation of 15 μM Aβ_40_ peptides, derived from fitting Eq. 1 to the ThT fluorescence curves shown in Fig. 2.

### AFM imaging

AFM images were recorded for some of the samples used in the ThT fluorescence measurements, i.e. the two control samples of 15 μM Aβ_40_ peptides in buffer with and without 2% DMSO, and the samples with additions of 15 μM, 75 μM, and 300 μM of 6-gingerol (Fig. 3). These samples were incubated for 72 h, to ensure aggregation into the mature elongated fibrils seen in Fig. 3A. Incubation in the presence of 2% DMSO produced similar fibrils, although together with small non-fibrillar clumps (Fig. 3B). Somewhat similar results, although with even more clumps, were obtained for the samples incubated together with 15 and 75 μM 6-gingerol, which also contained 0.1% and 0.6% DMSO, respectively (Figs. 3C and 3D). The sample with 300 μM of 6-gingerol and 2% DMSO does however display a different morphology, as it clearly contains more amorphous clumps than elongated fibrils (Fig. 3E). When evaluating these samples, it is a confounding factor that DMSO appears to slightly affect the fibril formation. The sample with 300 μM 6-gingerol however contains 2% DMSO (Fig. 3E), i.e. the same amount of DMSO as the control sample with DMSO (Fig. 3B). Thus, the different morphologies of the Aβ_40_ aggregates in these two samples is clearly caused by the added 6-gingerol and not by the DMSO alone.

**Figure 3.**
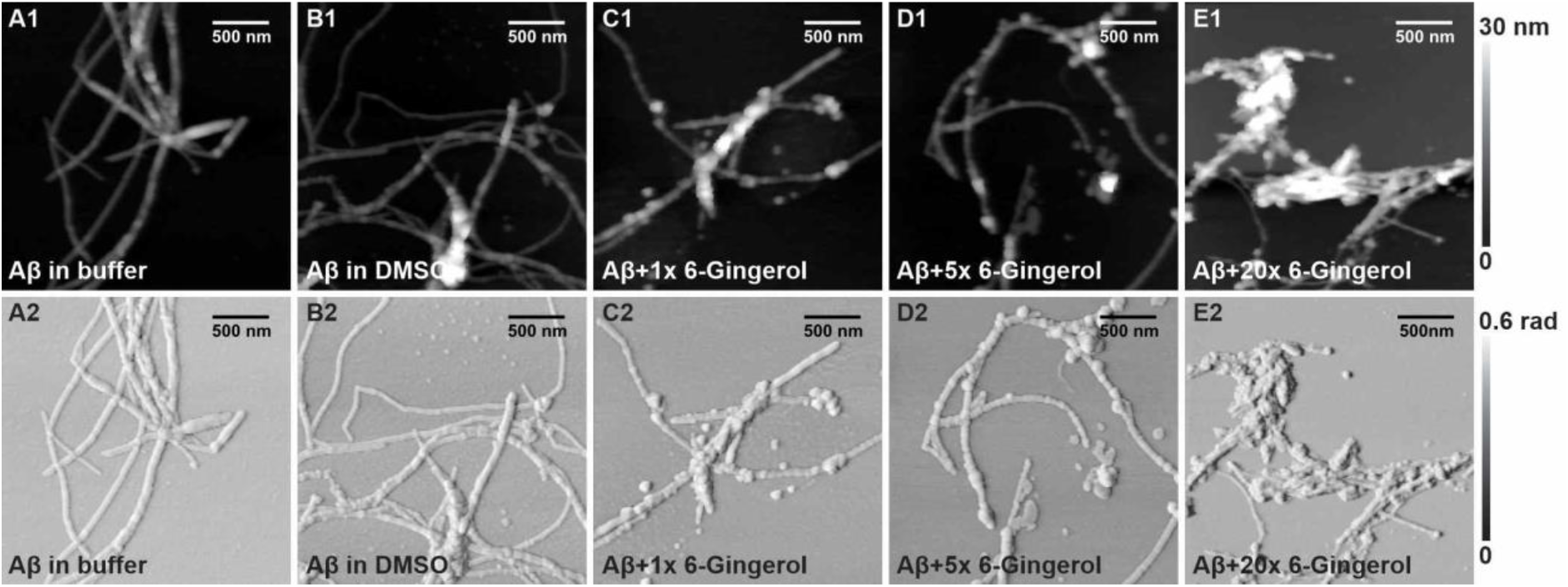
AFM images showing aggregates of 15 μM Aβ_40_ peptide. (A) Aβ_40_ in buffer. (B) Aβ_40_ in DMSO. (C) Aβ_40_ and 15 μM 6-gingerol in DMSO, (D) Aβ_40_ and 75 μM 6-gingerol in DMSO, (E) Aβ_40_ and 300 μM 6-gingerol in DMSO. Top row: height profiles. Bottom row: mechanical phase images.

### NMR spectroscopy

NMR experiments were conducted to investigate possible molecular interactions between 6-gingerol and the monomeric Aβ_40_ peptide. The finger-print region of the ^1^H,^15^N-HSQC spectrum of 92 μM monomeric ^15^N-labeled Aβ_40_ peptide is shown in Fig. 4 (blue spectrum), both for Aβ_40_ in buffer and for Aβ_40_ bound to SDS micelles. The SDS micelles were here used as a simple model for a membrane environment that is suitable for NMR studies (Österlund *et al.*, 2018a; Österlund *et al.*, 2018b). In both environments, addition of DMSO (2% in the buffer sample and 3% in the sample with SDS micelles) induces chemical shifts of most crosspeaks (Fig. 4, red spectra). This is consistent with previous NMR studies of Aβ_40_ in DMSO (Wallin *et al.*, 2017). Addition of 6-gingerol dissolved in DMSO increased the DMSO concentration to 4% in the buffer sample and to 5% in the sample with SDS micelles. This addition induces chemical shift changes for the NMR crosspeaks that are perfectly consistent with the changes induced by DMSO alone (Fig. 4, orange spectra). This shows that 6-gingerol does not have any strong interaction of its own with monomeric Aβ_40_, neither in aqueous solution nor in a membrane environment.

**Figure 4.**
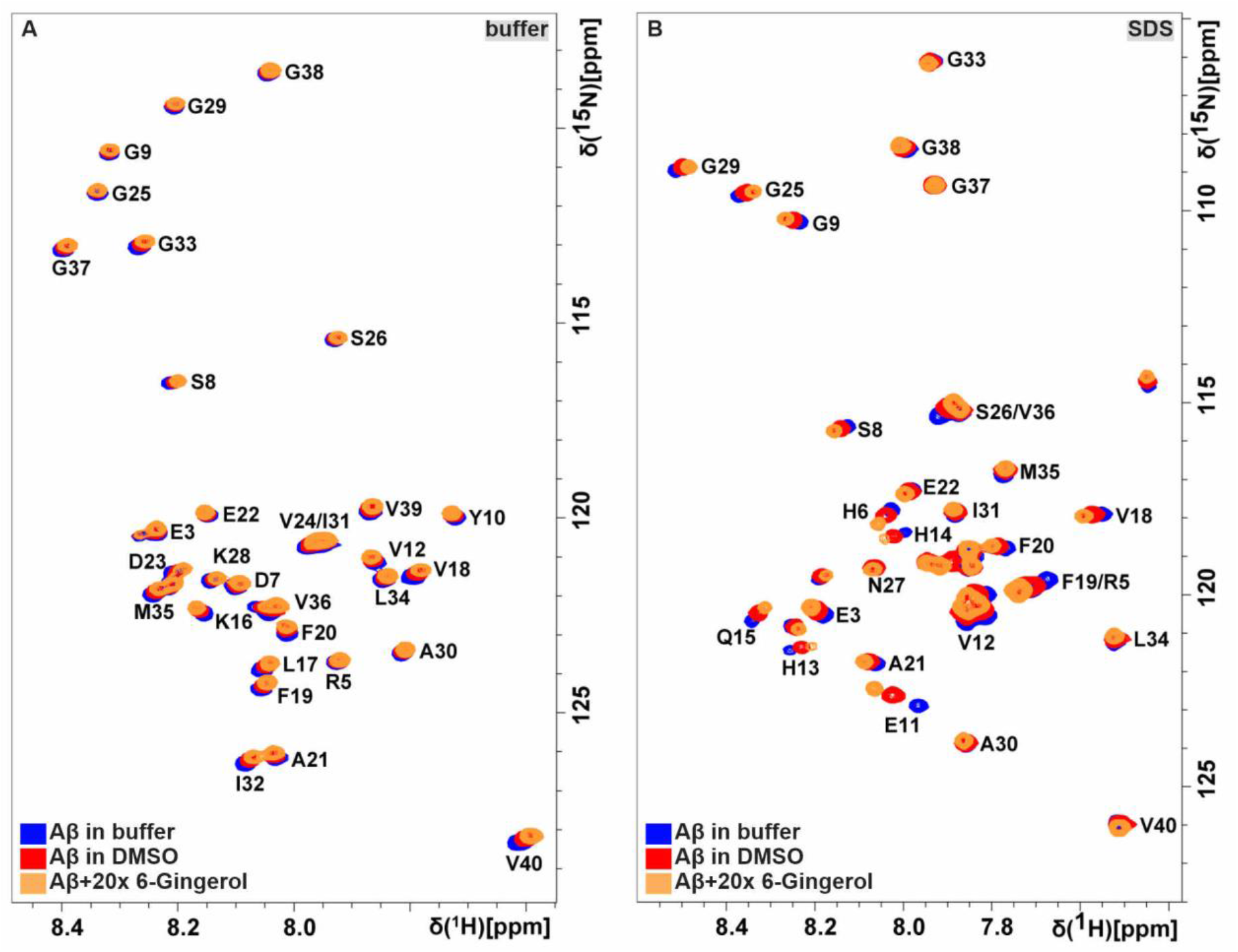
2D NMR ^1^H,^15^N-HSQC spectra recorded at +20 °C for 92 μM monomeric Aβ_40_ peptide in 20 mM sodium phosphate buffer, pH 7.3, for (A) Aβ_40_ in buffer alone, and (B) Aβ_40_ bound to micelles of 50 mM SDS. The spectra were recorded before (blue) and after addition of DMSO (red), and then after addition of 1.84 mM 6-gingerol in DMSO.

## DISCUSSION

Given the ancient history and cultural importance of ginger in many parts of the world (Iranshahy and Javadi, 2019; Khodaie and Sadeghpoor, 2015; Saenghong *et al.*, 2012), it is desirable to understand the molecular mechanisms behind its proposed benefits to human health. Such mechanistic investigations may also expand ethnomedical research, which often focuses on population-level medical effects and exposure/uptake levels (Sholts *et al.*, 2017; Wärmländer *et al.*, 2011).

Here, we show that 6-gingerol interferes with the aggregation mechanisms of Aβ_40_ peptide aggregation, by inducing aggregation into amorphous clumps rather than into elongated fibrils (Fig. 3). Our ThT fluorescence assays show that 6-gingerol has no systematic effect on the kinetics of the Aβ_40_ aggregation process, and that approximately the same amount of amyloid aggregates is formed with and without 6-gingerol (Fig. 2). From a medical perspective, however, the most important aspect of Aβ aggregation may not be the amount or speed of aggregation, but rather the properties of the aggregates. The neuronal death in AD appears to be mainly caused by small oligomeric Aβ aggregates of unknown composition and structure (Luo *et al.*, 2014b; Sardar Sinha *et al.*, 2018; Sengupta *et al.*, 2016) that might disrupt cell membranes (Wärmländer *et al.*, 2019). Thus, the observed interference of 6-gingerol with the Aβ aggregation processes could provide a molecular explanation of the previously observed beneficial effects of gingerols on cell and animal models of AD pathology (Choi *et al.*, 2018; Halawany *et al.*, 2017; Jeong *et al.*, 2013; Mohd Sahardi and Makpol, 2019; Wang *et al.*, 2014a; Zeng *et al.*, 2015).

The NMR results show that 6-gingerol does not interact with monomeric Aβ_40_, neither in aqueous solution nor in membrane-mimicking micelles. Thus, interaction appears to take place only when oligomers or larger aggregates have formed. This is not unreasonable, as Aβ oligomers are considered to be more hydrophobic than the amphiphilic Aβ monomers (Wärmländer *et al.*, 2019), and thus more likely to interact with the hydrophobic 6-gingerol molecules. In fact, the ideal AD drug is a molecule that interferes with toxic Aβ aggregates but not with the Aβ monomers, as the latter may have beneficial biological functions in their non-aggregated form (Dominy et al., 2019; Frozza et al., 2018; Querfurth and LaFerla, 2010; Rajendran and Annaert, 2012).

As a molecule that is non-toxic (Kaul and Joshi, 2001), easy to produce and administer, and small enough to easily pass through the blood-brain-barrier, 6-gingerol has suitable properties for use as a drug. This study suggests that 6-gingerol may be used to combat AD by interfering with the aggregation of Aβ peptides.

## ACKNOWLEDGMENTS

We thank Teodor Svantesson and Georgia Pilkington for helpful discussions and advice.

